# Human norovirus infection of primary B cells triggers immune activation *in vitro*

**DOI:** 10.1101/2021.05.14.444272

**Authors:** Carmen Mirabelli, Melissa K. Jones, Vivienne Young, Abimbola O. Kolawole, Irene Owusu, Mengrou Shan, Basel Abuaita, Irina Grigorova, Steven K. Lundy, Costas A. Lyssiotis, Vernon K. Ward, Stephanie M. Karst, Christiane E. Wobus

## Abstract

Human norovirus (HNoV) is a global health and socio-economic burden, estimated to infect every individual at least five times during their lifetime. The underlying mechanism for the potential lack of long-term immune protection from HNoV infections is not understood and prompted us to investigate HNoV susceptibility of primary human B cells and its functional impact. Primary B cells isolated from whole-blood were infected with HNoV-positive stool samples and harvested 3 days post infection (dpi) to assess viral RNA yield by RT-qPCR. A 3-18 fold increase in HNoV RNA yield was observed in 50-60% donors. Infection was further confirmed in B cells derived from splenic and lymph node biopsies. Next, we characterized infection of whole-blood derived B cells by flow cytometry in specific functional B cell subsets (naïve CD27^-^IgD^+^, memory switched CD27^+^IgD^-^, memory unswitched CD27^+^IgD^+^ and double-negative CD27^-^IgD^-^). While susceptibility of subsets was similar, we observed changes in B cell subsets distribution upon infection that were recapitulated after treatment with HNoV virus-like particles and mRNA encoding for HNoV NS1-2 protein. Importantly, treatment of immortalized BJAB B cell lines with the predicted recombinant NS1 protein triggered cell proliferation, increased ATP production, and induced metabolic changes, as detected by means of CFSE/Ki67 staining, seahorse analysis and metabolomics, respectively. These data demonstrate the susceptibility of primary B cells to HNoV infection and suggest that the secreted NS1 protein affects B cell function, proliferation and metabolism *in vitro*, which could have implications for viral pathogenesis and immune response *in vivo*.

**Importance:** Human norovirus (HNoV) is the most prevalent causative agent of gastroenteritis worldwide. Infection results in a self-limiting disease that can become chronic and severe in the immunocompromised, elderly and infants. There are currently no approved therapeutic and preventative strategies to limit the health and socio-economic burden associated with HNoV infections. Moreover, HNoV does not elicit life-long immunity as repeat infections are common, presenting a challenge for vaccine development. Given the importance of B cells for humoral immunity, we investigated susceptibility and impact of HNoV infection on human B cells. We found that HNoV replicates in human primary B cells derived from blood, spleen and lymph nodes specimens and induces functional changes in B cells, mediated in part by the non-structural protein NS1. Because of the secreted nature of NS1, we put forward the hypothesis that HNoV infection can modulate bystander B cell function with potential implications in systemic immune response.

## INTRODUCTION

Human norovirus (HNoV) is the most prevalent virus associated with foodborne illnesses, specifically viral gastroenteritis, that are considered by the World Health Organization as a major public health concern (1). In addition, the economic burden of HNoV worldwide has been estimated at 60 billion USD per year (2). Development of effective therapeutics has been hampered in part by the lack of a robust culturing system. Despite the ability of HNoV to infect human intestinal enteroids and immortalized B cells (3, 4), no cell-derived HNoV stock has been produced yet and infection is routinely performed with stool samples that are HNoV-positive by qPCR. The report of modest viral replication in a line of immortalized B cells, the BJAB (4), has opened a controversy in the field on whether or not B cells can support productive viral replication *in vivo* (5). Human norovirus antigen was detected in the lamina propria of both humans (in macrophages, dendritic cells and T cells of biopsies from two immunocompromised patients) (6) and animal models (in macrophages, lymphocytes, and dendritic cells of piglets; in dendritic cells of chimpanzee and cells of the hematopoietic lineage in zebrafish larvae) (7–9) but not specifically or exclusively in B cells. In addition, infection of common variable immunodeficiency-patients results in chronic infection and continuous symptomatology suggesting that immune cell infection is not absolutely required for HNoV susceptibility or induced pathophysiology *in vivo* (10).

On the other hand, it has been estimated that every person experiences HNoV at least five times in their lifetime suggesting a lack of long-term immune protection (11) but the underlying mechanisms are not understood. Broad protection and long-term duration of protection are also critical parameters in the development of an effective HNoV vaccine, which is lacking to date (12). B cells are a critical component of effective, long-term immunity. There is therefore a need for targeted studies that explore the relationship between HNoV infection and B cells. In this manuscript, we sought to determine whether primary human B cell support *ex vivo* infection with HNoV. Increases in HNoV genome levels were observed in primary human B cells derived from blood, spleen and lymph nodes and infection was blocked with either the nucleoside inhibitor 2’CMC or type I interferons. In addition, infection with HNoV but also treatment with HNoV virus-like particle (VLPs) or with the non-structural protein NS1 affected B cells functional subsets distribution over time. Previous work demonstrated that the NS1 protein is secreted during infection with murine and human NoV (13). Our study now discovered that treatment with the predicted NS1 protein alone induces B cell proliferation and increases glucose metabolism and OXPHOS, suggesting that this viral protein can induce bystander B cell activation. The implications of this finding on HNoV immune responses are potentially manifold and call for more detailed *in vivo* studies.

## RESULTS

### HNoV replicates in primary B cells *in vitro* and is restricted by the type I interferon response

To determine whether primary human B cells were susceptible to HNoV infection, peripheral blood mononuclear cells (PBMCs) were obtained from the blood of different unidentified donors after ficoll centrifugation and B cells were isolated by using magnetic beads coupled to anti-CD19. The B cells were co-cultured with γ-irradiated human CD40 ligand (hCD40L)-expressing 3T3 cells for 2 days and subsequently infected with HNoV-positive stool samples of GII.4 or GII.6 genotypes. At 3 days post infection (dpi), viral RNA was measured by reverse transcriptase (RT)-qPCR and increase in viral replication was calculated as a fold increase (FI) *vs* 0 dpi (inoculum). Primary B cells derived from 6/12 (50%) and 11/18 (60%) donors were permissive to HNoV GII.4 and GII.6 replication, respectively, with FI ≥ than 3 (**Figure 1A**). A threshold of 3-FI was previously defined as indicator of HNoV replication *in vitro* (14). Importantly, upon treatment with the nucleoside analogue 2’CMC, an *in vitro* inhibitor of NoV replication, no increase of viral RNA was detected in any of the donors tested (**Figure 1B**), suggesting that HNoV actively replicate in primary B cells. To confirm this finding, primary B cells were isolated from human spleen and lymph node biopsies with EasySep™ Human B Cell Isolation Kit (StemCell technologies) and were infected with GII.6 HNoV-positive stool samples. BJAB, a clone of immortalized B cells, that was previously described to support HNoV infection (4) served as a control. Both B cells from spleen and lymph node supported HNoV replication, with 80% (8/10) of donors permissive to infection in the case of lymph node B cells and 100% (6/6) of donors for splenic B cells (**Figure 1C**). Interestingly, splenic and lymph node B cells were more permissive to infection than blood-derived B cells, regardless the genetic background of the different donors (**compare Figure 1C vs 1A**). Since HNoV infection of intestinal epithelial cells is restricted by interferons (IFNs) (15), we next tested whether HNoV infection of primary B cells was similarly susceptible to type I IFNs. Primary splenic B cells and BJAB were treated with IFNβ (1000 U/mL) for 24 hrs prior to infection. The treatment reduced infection of primary B cells by 8-fold but only 3-fold in BJAB, suggesting that primary B cells are more sensitive to IFN treatment than BJAB (**Figure 1D**). Conversely, when primary B cells from spleen tissues were pretreated for 18 hrs with antibodies neutralizing IFNα (1:4000), IFNβ (1:4000), IFNβ2 (1:4000), the type I IFNαβ receptor (1:1000), or a combination of antibodies (at the concentrations above), HNoV infection increased at least 2-fold (**Figure 1E**). Together, these data suggest that primary B cells can support HNoV replication *ex vivo*, albeit modestly, and that infection is sensitive to the nucleoside analog 2’CMC and the antiviral activity of type I IFNs.

**Fig 1.**
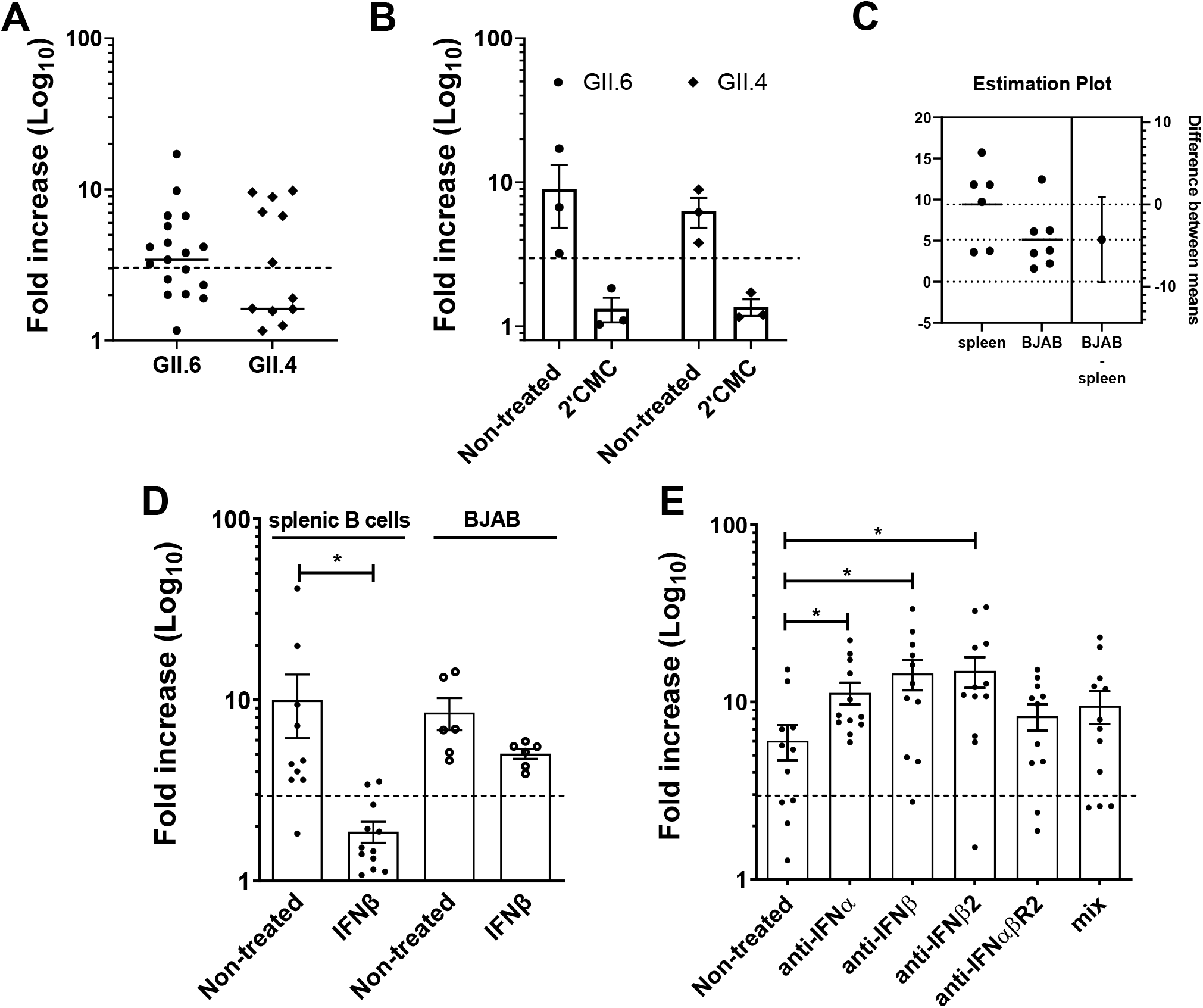
HNoV replicates in primary B cells and replication is restricted by interferon. **A)** Primary B cells were extracted from whole blood and infected with GII.4 and GII.6 positive stool sample in 1.5ml tubes. One hour after adsorption, cells were washed and incubated for 3 days at 37°C in co-culture with hCD40L-3T3. Graph represents the fold increase in viral genome copies obtained by RT-qPCR of 3 dpi *vs* 0 dpi. Each dataspoint represent a biological replicate (different blood donor) **B)** Primary B cells were infected with GII.4 and GII.6 positive stool sample in the presence of 2’methyl cytidine (2’CMC). Graph represent fold increase in viral genome copies. Each dot is a single biological replicate. **C)** Primary B cells were isolated from spleen and lymph nodes of patients’ biopsies. The graph represents the fold increase in viral genome copies. Each dot is a single biological replicate (single donor), that is average of three technical replicates **D)** Primary B cells from spleen and BJABs were treated with IFNβ prior to infection with HNoV-positive stool samples. **F)** Primary B cells from spleen were treated with anti-IFN antibodies prior to and through the duration of infection with HNoV-positive stool. For D and F, each dot represents a technical replicate of infection. The dashed line represents the threshold of 3-FI, indicative of viral replication.

### HNoV infection efficacy of blood-derived B cells is dependent on donor and culturing time

To test whether the efficacy of HNoV infection could be improved in primary B cells, B cells were isolated from whole blood of different donors and infected directly after isolation or co-cultured with γ-irradiated human CD40 ligand (hCD40L)-expressing 3T3 cells for 2 and 5 days before infection. Infection with GII.6-positive stool samples was more efficient in freshly isolated B cells as compared to primary B cells in culture for day 2 or day 5, although cell viability did not change over time (**Figure 2A**). Even in HNoV-infected B cells isolated from the same donor, infection efficiency decreased over time of culture (**Supplementary figure 1A**). In order to characterize HNoV infection in primary B cell, a flow cytometry pipeline was established to identify functional B cells subsets according to the CD27 marker of memory and IgD expression on the cell surface; naïve (CD27^-^IgD^+^), memory switched (CD27^+^IgD^-^), memory unswitched (CD27^+^IgD^+^) and double-negative (CD27^-^IgD^-^) B cells (**Supplementary figure 1B**). The prevalence of each subset was first determined in non-infected primary B cells freshly isolated or cultured on hCD40L-3T3 cells for 2 and 5 days. A change in subset distribution was consistently observed across donors (**Figure 2D**). The concomitant decrease of HNoV replication and loss of specific B cell subsets over time raised the possibility that HNoV selectively infects specific functional B cell subset(s).

**Fig 2.**
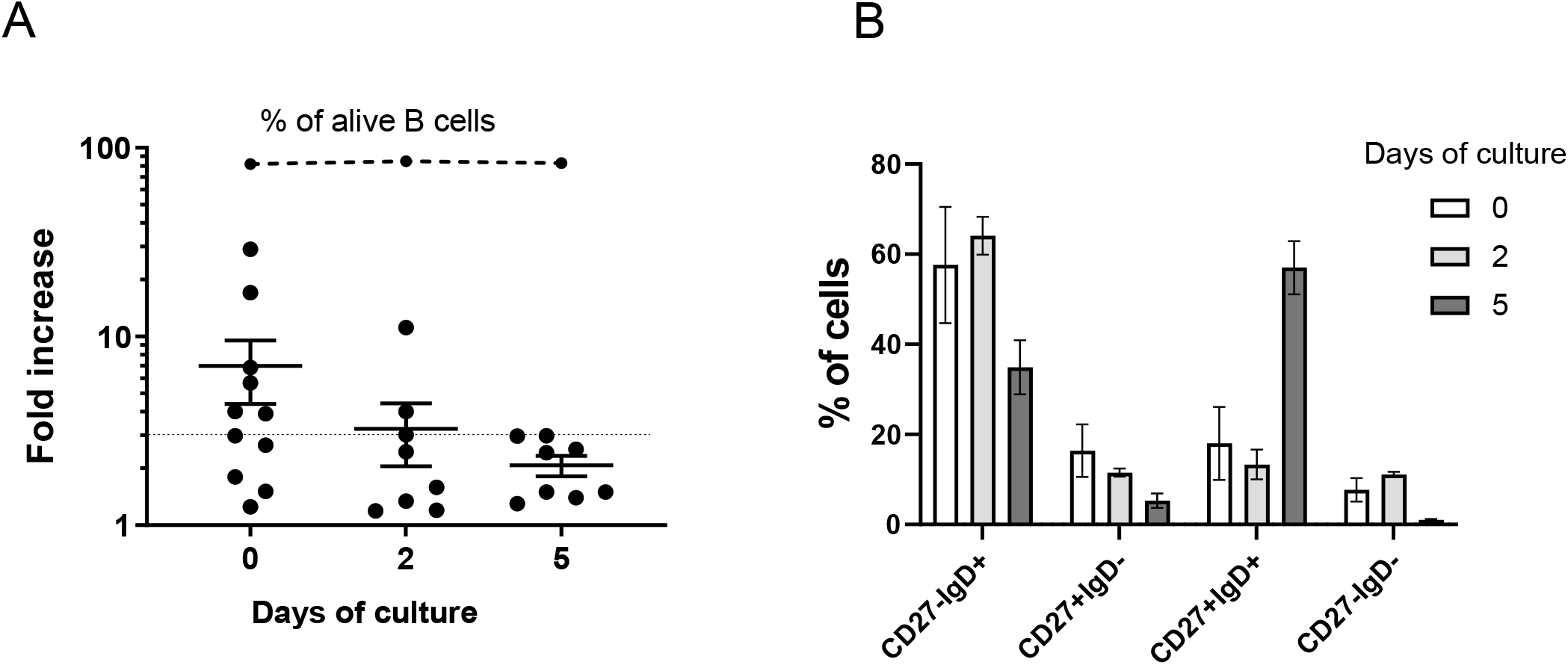
HNoV infects more efficiently freshly isolated B cells. **A)** Whole-blood derived B cells were infected with GII.6 HNoV-positive stool sample at the moment of isolation or after 2 or 5 days of co-culture with hCD40L-3T3 cells. Graph represents the fold increase in viral genome copies obtained by RT-qPCR of 3dpi *vs* day 0 dpi. Cell viability was determined on uninfected cells by flow cytometry, as a percentage of Live/dead negative after staining with LIVE/DEAD Fixable Aqua Dead Cell Stain. Each data-point in the graph indicates a biological replicate from one donor (day 0, N=11, day 2 and day 5, N=8). Statistics were run on GraphPad Prism and the day 5 group is statistically different then day 0 group by Mann-Whitney test, p=0.035 **B)** Primary B cells from various donors were subjected to flow cytometry analysis after isolation (day 0, N=10) and at day 2 (N=5) and day 5 (N=5) of co-culture with hCD40L-3T3 cells. Percentage of CD27^-^ IgD^+^ (naive), CD27^+^IgD^-^ (memory switched), CD27^+^IgD^+^ (memory unswitched) and CD27^-^ IgD^-^ (double-negative) was calculated with FlowJo software and represent average, SD of 5-10 independent experiments.

### HNoV tropism is not restricted in a specific B cell subset

To test this hypothesis, freshly isolated blood-derived primary B cells were infected with HNoV GII.6-positive stool and were subjected to flow cytometry analysis at 3 dpi. Cells replicating HNoV genome were detected by using an antibody against double-stranded (ds) RNA, an intermediate of viral replication. HNoV-infected cells ranged between 5-10% of total B cells in donors that were permissive to infection by RT-qPCR (FI>3) whereas a lower percentage of infected cells corresponded to non-permissive donors (FI<3) (**Figure 3A**). A representative flow plot of one permissive and one non-permissive donor is shown in **Figure 3B**. However, the proportion of HNoV-infected cells did not differ across the functional B cell subsets (**Figure 3C**), suggesting that HNoV tropism is not restricted to specific B cell subset(s).

**Fig. 3.**
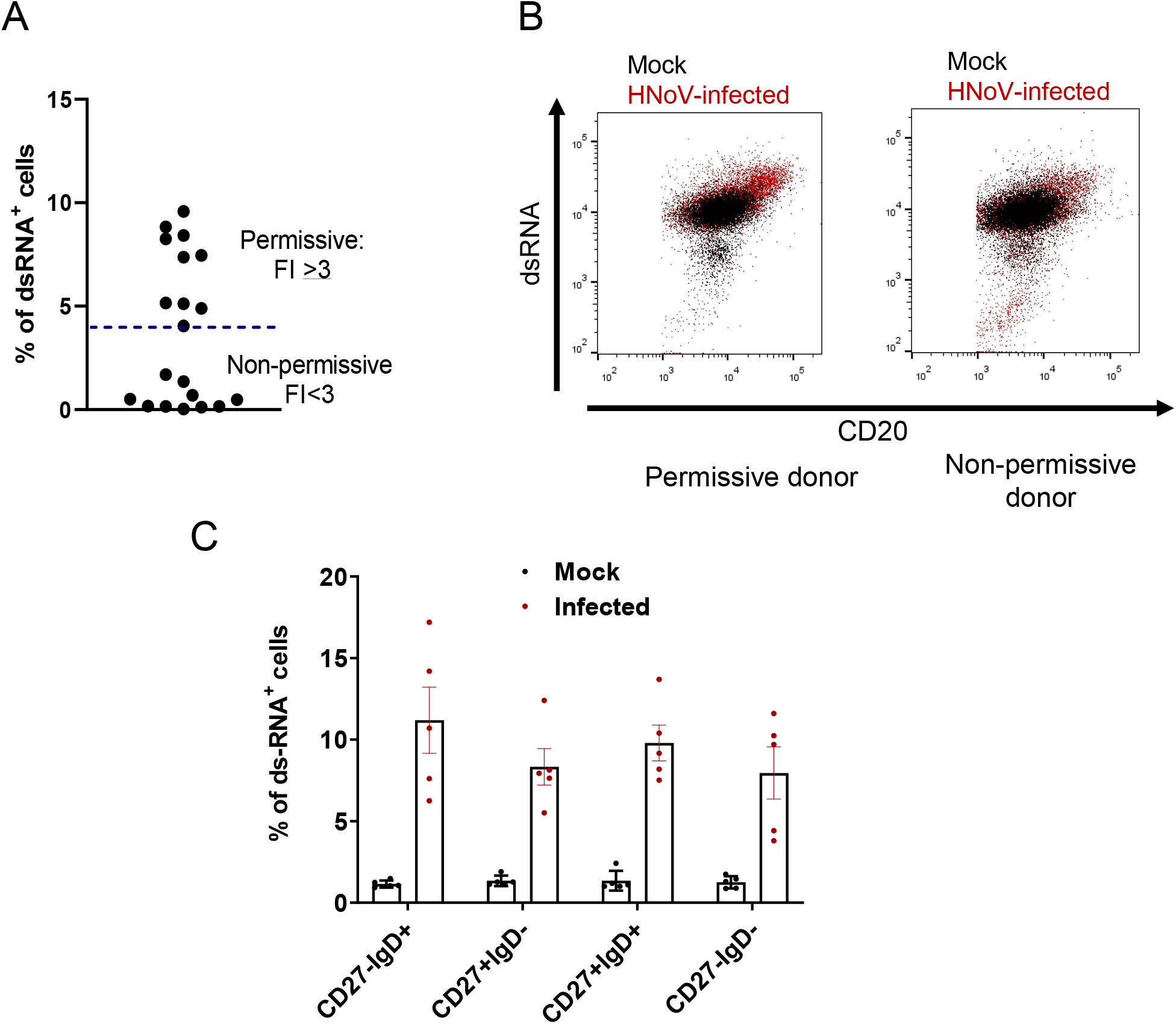
HNoV tropism is not restricted to a specific B cell subset. **A)** Freshly isolated primary B cells were infected or mock-infected with GII.6 positive stool sample and harvested at 3 dpi for flow cytometry analysis after staining with LIVE/DEAD Fixable Aqua Dead Cell Stain, PerCP/Cy5.5 anti-human CD20, PE-CF594 Mouse Anti-Human CD27, Pacific Blue™ anti-human IgD and biotynilated-dsRNA (J2) for the detection of infected cells. B cells were also infected in parallel for to determine fold increase (FI) of viral replication by RT-qPCR. The graph shows the percentage of ds-RNA^+^ cells of infected B cells. Each data point is an independent biological replication from a different donor. Permissive donors were defined according to 3-FI or greater as defined by RT-qPCR. **B)** Representative flow plots of infected and mock-infected permissive and non-permissive donors. **C)** Percentage of ds-RNA^+^ cells in the different functional B cell subsets. The dsRNA^+^ gate was arbitrarily defined on the HNoV-infected non-permissive donors to include ≤1% event. Percentage are calculated in the infected gate (4-10% of total B cell). Each datapoint on the graph represents an independent biological replicate of N=5 different permissive donors. Average and SD are also depicted.

### HNoV infection or treatment with selected viral proteins induces changes in functional B cell subset distribution

To assess the potential impact of HNoV on B cell function, we first compared the distribution of B cell subsets between permissive and non-permissive donors upon infection. Briefly, from the % of cells in each subset, we calculated the ratio in the HNoV-infected *vs* Mock (non-infected) condition, whereby ratio of 1 represents no changes, ratio >1 represents an increase and ratio <1 represents a decrease in subset distribution, respectively. We found that upon HNoV-infection, double-negative (CD27^-^IgD^-^) and unswitched (CD27^+^IgD^+^) B cells exhibited bi-modal distributions in permissive donors but no changes were observed in non-permissive donors (**Figure 4A**). As a technical control for the flow cytometry pipeline and a biological control to determine the extent of these changes in subset distribution, we treated primary B cells with interleukin (IL) 4 (20 ng/ml), a cytokine known to promote switching *in vitro* (16). As expected, IL-4 treatment resulted in an enrichment in switched and double-negative (IgD^-^) subsets (**Figure 4B**) and the magnitude of the changes was comparable to the levels observed during infection (**Figure 4A**). To define the molecular triggers for the HNoV-induced changes, primary B cells from different donors were treated with GII.4 HNoV VLPs (1 μg/ml) or transfected with the synthetic dsRNA mimic poly (I:C) that serves as a control for viral replication, or with 1 µg of mRNA encoding HNoV NS1-2 protein. We used transfection reagent alone as a control for the poly (I:C) and NS1-transfected condition and analyzed B cell sub-population distribution at 3 days post-treatment/transfection (dpt), consistently with the infection timeframe. Treatment with HNoV VLPs and transfection of NS1-2 mRNA, but not poly I:C, induced changes in subsets distribution (**Figure 4C**). In particular, expression of NS1-2 induced bi-modal distribution changes reminiscent of those induced by HNoV infection (**compare Figure 4A vs. C**). Together, these data highlight the impact of viral infection and viral proteins on functional B cell subset distribution. In addition, given the previously reported secreted nature of the non-structural protein NS1 (13), these data suggest that NS1 may alter bystander B cell function.

**Fig. 4.**
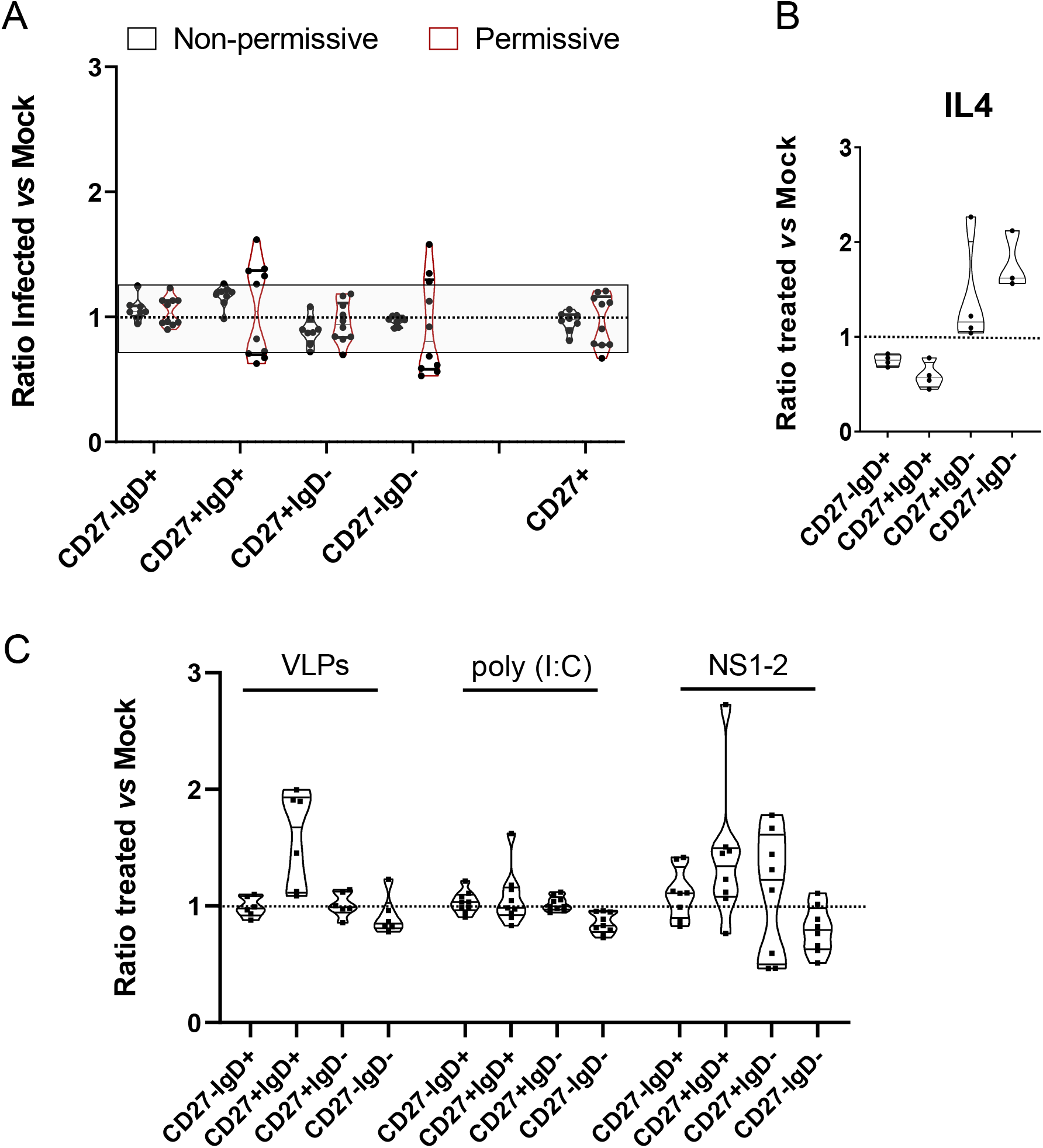
HNoV infection or treatment with selected viral proteins induce changes in B cell functional subsets distribution. **A)** Changes in B cell subsets expressed as a ratio of percentages in HNoV-infected *vs* non-infected (Mock). Permissive (N=10) and non-permissive (N=8) donors were defined according to FI of viral replication. Values close to 1 represent no or little changes, values >1 represents increase and values <1 represents decrease in percentage of cells for each sub-population, respectively. **B)** Changes in B cell subsets upon treatment with IL4 (20ng/ml) for 3 days, expressed as a ratio of treated *vs* non-treated (Mock). **C)** Changes in B cell subpopulation 3dpt with HNoV virus-like particles (VLPs, 1µg/ml), Poly (I:C) and HNoV NS1-2 mRNA (1µg), expressed as a ratio of treated *vs* non-treated (Mock). Violin plots depicts mean, interquartile and each dot represents an independent biological replicate (single donor).

### NS1 protein increases proliferation and metabolism of an immortalized B cell line

To characterize the changes triggered by the viral protein NS1 in B cells, the immortalized B cell line BJAB was used as a simplified homogeneous cell culture system. To mimic the secreted nature of NS1, we produced a recombinant, baculovirus-expressed HNoV NS1 protein that encompasses the disordered region (1-134 aa) from the GII.4 Sydney 2012 strain (**Supplementary figure 2**). We first treated BJAB with 10 μg/ml of the putative HNoV NS1 protein and evaluated cell proliferation after CFSE and Ki67 staining at 3 dpt by flow cytometry. NS1 correlated with a lower CFSE signal and increased Ki67 staining that was 2 - 10 times higher than non-treated BJAB cells (**Figure 5A and 5B, Supplementary figure 3A and 3B**). B cell proliferation is typically associated with B cell activation, of which an increase in glucose uptake, glucose metabolism and oxidative phosphorylation are hallmarks (17). Thus, we next determined the effect of NS1 treatment on BJAB metabolism. We first analyzed the transcriptional activation of two genes involved in glucose metabolism, the glucose transporter GLUT-1 and hexokinase HK-1, which phosphorylates glucose to produce glucose-6-phosphate, the first step in most glucose metabolic pathways. At 3 dpt with 10 µg/ml of NS1, RNA was isolated and analyzed by qPCR. GLUT-1 mRNA levels did not significantly change, whereas HK-1 mRNA expression significantly increased (**Figure 5C**). Next, we performed a treatment of BJAB cells for 16 hrs with 1 µg/ml NS1 and analyzed the intracellular ATP levels by Seahorse. NS1 protein inactivated at 95°C for 10 minutes was used as a negative control, and cells treated with 1 µg/ml LPS were used as a control for B cell activation and metabolic reprogramming (18). NS1 protein treatment correlated with increased levels of oxygen consumption rate (OCR) that is a measure of mitochondrial oxidative phosphorylation while no significant changes were observed for extracellular acidification rate (ECAR) that is a measure of glycolysis (**Figure 5D**). These data were consistent with previous findings on B cells activation *in vitro* (17). Last, we confirmed these metabolic changes by performing liquid chromatography coupled tandem mass spectrometry (LC/MS)-based metabolomics analysis (19) of BJAB after 16 hrs stimulation with 1 µg/ml of NS1 (**Figure 5E**). Mannose-1 phosphate and fructose-6 phosphate were two of the most significantly altered metabolites. The modest increase of lactate and the high levels of fructose-6-phosphates and mannose are consistent with published data on upregulation of the glucose metabolism during B cell activation (18). Together, these data suggest that the viral protein NS1 induces metabolic changes in B cells that might lead to their proliferation and activation.

**Fig. 5.**
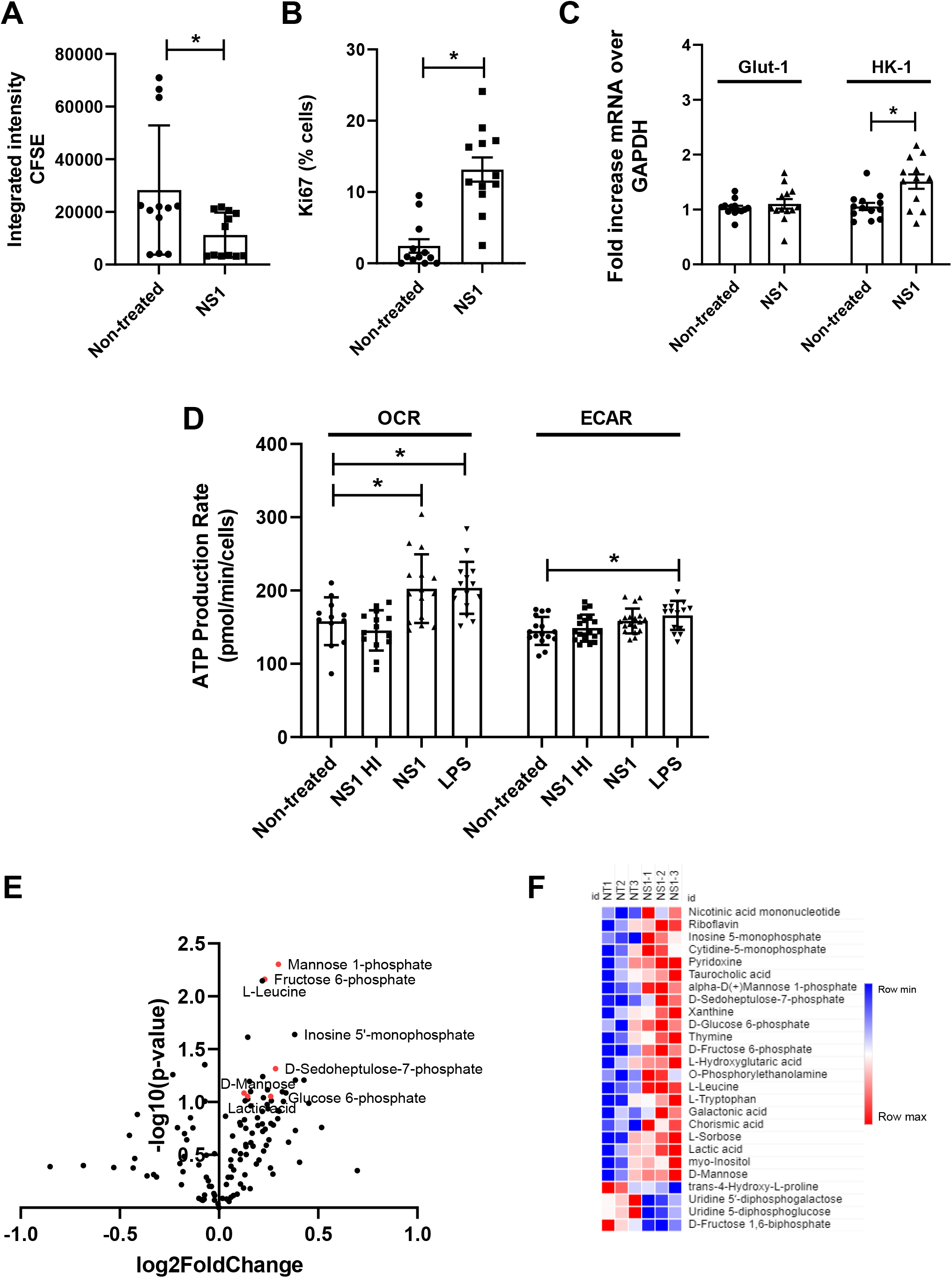
NS1 treatment enhances BJAB proliferation and metabolism. **A, B)** BJAB cells were stained with CFSE and treated (or mock-treated) with HNoV NS1 protein (10µg/ml) for 3 days. BJAB were harvested by fixation/permeabilization and stained with Ki-67. Cells were analyzed by flow cytometry with BD Fortessa and FlowJo software. Each data point represents a technical replicate from N=4 independent biological experiments. **C)** BJAB cells were treated with HNoV NS1 protein (10µg/ml) for 3 days. GLUT-1 and HK-1 mRNA were quantified by qPCR. GAPDH was used as an internal control. Each data point represents the mRNA fold increase over non-treated BJAB and it is a technical replicate from N=4 independent biological experiments. **D)** BJAB cells were treated with HNoV NS1 protein (1µg/ml), NS1 protein inactivated at 95°C for 10 minutes and LPS (1µg/ml) for 16hrs. 5×10^4^ BJAB cells/well were transferred in a Seahorse 96-well plate, washed with assay medium twice and kept in a non-CO_2_ incubator for 40 minutes. Cellular ATP was quantified with Seahorse XFe96 Extracellular Flux Analyzer. Data were plotted with Agilent ATP Assay Report Generator. Graphs represents average + SD of N=5 biological replicates (with at least 3 technical replicate, each). OCR= oxygen consumption rate (mitochondrial OXPHOS), ECAR= extracellular acidification rate (glycolysis). **E-F)** 10^6^ BJAB cells were treated with HNoV NS1 protein (1µg/ml) for 16hrs. Cells were harvested and metabolites were collected in 80% ice-cold ethanol followed by snapshot metabolomics analysis. **E**) Volcano plot where x-axis represents log2FoldChange of NS1 treated *vs* NT control cells and y-axis represents –log10 of the p-value. **F**) Heat-map of metabolites significantly upregulated (red) and downregulated (blue) in the NS1 condition *vs* non-treated. The three technical replicates are shown in the heat-map.

## DISCUSSION

In this study, we sought to determine whether human primary B cells are susceptible of HNoV infection *ex vivo* and whether infection has an impact on B cells function. We first demonstrated that HNoV actively replicates in human B cells, derived either from whole blood PBMCs or from patient’s spleen and lymph nodes biopsies. Treatment with the nucleoside 2’CMC or type I IFN abolished HNoV replication, suggesting that B cells are susceptible and permissive to infection. However, given the low level of increase in viral genome titers or dsRNA, it is possible that HNoV replication in B cells is defective and can result in abortive infection. We established that B cells derived from 80-100% of spleen and lymph nodes but only 50-60% of blood donors were permissive for HNoV replication pointing towards donor-specific determinants of permissiveness in the latter group. Future studies using paired samples from tissue and blood are needed to determine whether the susceptibility of B cells from different sites is similar or divergent. In addition, amongst the permissive blood donors, only 5-10% of primary blood-derived B cells were positive for viral dsRNA, suggesting that replication of HNoV is limited to a subset of B cells. We could not define this subset based on the memory marker CD27 and IgD, and further studies should be designed to define this population after FACS sorting, possibly by means of RNAseq or cytometry by time of flight (CyTOF). The low percentage of infected cells in the more homogeneous BJAB model (20) and also in the context of murine norovirus infection in B cells (7-9% of primary murine B cells [unpublished data] and ∼10% of M12 cell line (4)) suggests that other factors might define the susceptible subset (such as the expression levels of the viral receptor, the metabolic state of the cells, etc). Histo-blood group antigens have been identified as an attachment factor and genetic susceptibility factor for HNoV infection in the intestine (21). However, analysis of FUT2 expression, the enzyme required for addition of terminal fucose to carbohydrate chains, in a subset of B cell donors did not correlate with susceptibility (data not shown). Thus, the specific nature of the susceptibility factor(s) in B cells remains to be identified in future studies.

The second main finding in our manuscript was the ability of HNoV infection or treatment with viral proteins, specifically NS1, to drive changes in B cells functional subsets. Interestingly, we did not observe a specific enrichment or depletion of B cell subsets upon NS1 treatment, suggesting that additional stimuli are required to engage the B cell receptor (BCR) towards the differentiation into a functional subset, whereas stimulation with VLPs (via multivalent engagement of BCR) resulted in a specific increase of unswitched cells. In the more homogeneous model of BJAB B cells, we further demonstrated that NS1 treatment alone increases cell proliferation and cellular metabolism, features that are consistent with B cell activation. Whether replication of HNoV in B cell occurs *in vivo* remains unresolved. However, secretion of NS1 from other infected cell types could stimulate proliferation and metabolism of surrounding B cells. These data suggest a model whereby an enteric virus, through direct infection or bystander effects, induces changes in the activation profile of B cells and possibly other immune cells. Activation of B cells in the lamina propria or gut-associated lymphoid tissue (GALT) could have profound implications for the development of adaptive immune responses and protection from re-infection. Future studies will be required to test this hypothesis. In conclusion, our study demonstrates that a proportion of primary B cells is susceptible to HNoV infection *in vitro* and highlight a new function for NS1 in B cell activation with implications for viral pathogenesis.

## MATERIAL & METHODS

### Cell lines, compounds, virus

BJAB, a EBV-negative transformed B cell line was maintained in RPMI medium supplemented with 10% FBS, 1 mM L-glutamine and 1 mM penicillin/streptomycin at 37°C in a 5% CO_2_ incubator. 2′-*C*-Methylcytidine (2’CMC), an *in vitro* inhibitor of HNoV replication was supplied from Sigma-Aldrich. Human IFNβ was purchased from PBL Assay Science and used at a concentration of 1000 U/ml. IFN neutralizing antibodies were obtained from PBL Assay Sciences and were used at the following concentrations: anti-IFNα 1:4000), anti-IFNβ (1:4000), anti-IFNβ2 (1:4000), and anti-IFNαβR2 (1:1000). HNoV GII.4 virus-like particles (VLPs) were purchased from The Native Antigen Company, Poly (I:C) was obtained from Invivogen (tlrl-picwlv). Generation of the plasmid encoding HNoV GII.4 Sydney NS1-2 full-length protein was previously described (22). GII.4-Sydney and GII.6 HuNoV-positive stool samples were kindly provided by Dr. Vinje (Centers for Disease Control and Prevention, USA) and Dr. Karst (University of Florida, USA), respectively. Stool samples were diluted with PBS to make a 10% (w/v) stock solution. Diluted stool samples were vortexed and centrifuged at 20,000 x g for 1 min. Supernatants were used for infection.

### Primary B cell isolation from whole blood

30 ml PBS flow-through plasmapheresis filters or peripheral blood diluted in PBS (1:3) were collected in 50 ml tubes and underlaid with Ficoll-Paque (GE Healthcare). PBMC-containing buffy coat was obtained after centrifugation for 30 min at room temperature at 1200 RCF. After washing, PBMC were incubated with anti-CD19 magnetic beads (MACS, Miltenyi Biotech) in MagSep buffer (PBS-/-supplemented with 0.5% BSA and 2mM EDTA) for 15 min and washed in MagSep buffer twice before separation by flow on a magnetic column. The B cell fraction obtained after flow through was resuspended in maintenance buffer (IMDM supplemented with 10% fetal calf serum, 50ug/ml transferrin, 5ug/ml mixture of Transferrin/Insulin/Selenium, 15ug/ml of Gentamicin). Cells were cultured in the presence of previously ɣ-irradiated hCD40L-3T3 cells (23) at a ratio of 1:8.

### Infection of blood-derived primary B cells with HNoV

Freshly isolated B cells, or B cells that were co-cultured with hCD40L-3T3 cells for 2 or 5 days post isolation were infected with HNoV-positive stool samples of genotype GII.4 or GII.6. Briefly, for freshly isolated B cells, infection occurred in 1.5 ml tubes for 2h at 37°C followed by two washes with maintenance medium and seeding in the presence of ɣ-irradiated hCD40L-3T3 cells. For B cells in culture, HNoV-positive stool samples of genotype GII.4 or GII.6 was spin inoculated for 30 min at 800 x g at room temperature followed by two washing with maintenance medium. One batch of B cells was then harvested immediately after infection (day 0 of infection) with TRI Reagent (Zymo Research) and the rest of the infected cells were kept in culture for 3 days at 37°C. Infection was determined by RT-qPCR as the fold increase of viral genome copies at day 3 vs day 0 of infection.

### Ex-vivo isolation of splenic and lymph node B cell

Spleen and lymph node tissue samples were collected in compliance with the University of Florida Institutional Review Boards (IRBs) and protection of human subjects. De-identified biopsies were obtained and mashed in 100 µm and 70 µm cell strainers inside a Petri dish with 2 ml of PBS supplemented with 2% FBS. Cells were transferred into a 15 ml conical tube and centrifuged for 5 min at 500 x g. Germinal cells were resuspended in fresh media for counting while splenocytes were incubated in 1ml of ACK lysis buffer to remove red blood cells (ThermoFisher A1049201) for 3 min at room temperature and, after centrifugation, were resuspended in fresh media for counting. After counting, cells were centrifuged at 500 x g for 15 min at room temperature and were resuspended in RoboSep buffer for B cells isolation by negative selection with EasySep™ Human B Cell Isolation Kit (Stemcell Technologies) according to manufacturer’s instructions. After isolation, B cells were kept in complete culture medium (RPMI containing 10% FBS, Omega Scientific and supplemented with 1x Pen/Strep, Cellgro), seeded into 48-well plates at a concentration of 1 x 10^5^ cells per well and incubated 37°C at 5% CO_2_ overnight to 24 hrs prior to infection.

### Infection of germinal or splenic primary B cells with HNoV

HNoV-positive stool sample were diluted 1:10 in complete culture medium, and 100µl of virus prep was used to infect cells for 2hrs at 37° Cat 5% CO_2_. After infection, cells were centrifuged at 750 x g for 7.5 min and resuspended in culture media. Wells for day 0 of infection were immediately harvested in TRI Reagent (Zymo research) for RNA extraction. The remaining wells were incubated at 37°C for 3 days. For treatment with Type I IFN, immediately after plating, IFN was added to the wells and cells were incubated for 24 hrs prior to HNoV infection. For treatment with anti-IFN antibodies, antibodies were added to wells immediately after plating and incubated with cells for 18 hrs prior to infection. Media supplemented with anti-IFN antibodies was used throughout the infection and medium with antibodies were refreshed daily. Infection was determined by RT-qPCR as the fold increase of viral genome copies at day 3 vs day 0 of infection.

### Viral quantification by RT-qPCR and host genes qPCR

Viral RNA extraction was performed with the Direct-zol RNA MiniPrep Plus (Zymogen Research) according to the manufacturer’s protocol. HuNoV titers were determined by one step RT-qPCR as previously described (24). Host gene GLUT-1 and HK-1 transcripts were quantified by two step qPCR and normalized to glyceraldehyde 3-phosphate dehydrogenase (GAPDH) with the following primer sets (GLUT-1 F-TCAACACGGCCTTCACTG; GLUT-1 R-CACGATGCTCAGATAGGACATC; HK-1 F-GCACGATGTTCTCTGGGGTG; HK-1 R-CGTCAAGATGCTGCCAACCT; GAPDH F-TGGTTTGACAATGAATACGGCTAC, GAPDH R-GGTGGGTGGTCCAAGGTTTC).

### Flow cytometry analysis for B cell functional subsets

HNoV-infected or mock-infected primary B cells were harvested at selected times post infection in MagSep buffer (PBS supplemented with 0.5% BSA, 2mM EDTA). Cells were first stained with LIVE/DEAD Fixable Aqua Dead Cell Stain (Thermo Fisher Scientific) and after washing incubated with surface markers PerCP/Cy5.5 anti-human CD20 (BioLegend, 302326), PE-CF594 Mouse Anti-Human CD27 (BD BioScience, 562297) and Pacific Blue™ anti-human IgD (BioLegend, 348224). After 20 min, cells were washed and fixed/permeabilized with the Fixation/Permeabilization Solution Kit (BD Bioscience, 554714) according to manufacturer’s instructions. Next, cells were stained with the anti-double-stranded (ds) RNA antibody (J2, Scicons) -previously biotinylated with EZ-Link™ Micro NHS-PEG4-Biotinylation Kit (ThermoFisher 21955) to increase specificity and sensitivity of the assay- and with the APC/Cy7 streptavidin antibody (BioLegend, 405208). Data was acquired with BD Fortessa and analyzed by using FlowJo. Compensation was performed on uninfected BJAB.

### NS1 protein purification

The putative NS1 region (1-134aa) of the HNoV NS1 from the GII.4 Sydney 2012 strain (Accession no. JX459908.1) was expressed in *Trichoplusia ni* insect cells using the commercial recombinant baculovirus system, Flashback Ultra (Oxford Expression Technologies). The expression construct contained a N-terminal His-Strep II tag, a spidroin NT* solubility tag (25), an enterokinase cleavage site and a flexible linker GGSRS adjacent to HNoV NS1 (**Supplementary figure 2**). Following expression at 27°C for 3 days, the cells were lysed in 50 mM NaH_2_PO_4_.2H_2_O pH 8, 300 mM NaCl, 10% glycerol buffer containing 1% Triton X-100. The protein was purified on Streptactin XT superflow beads (IBA Lifesciences) and eluted with 50 mM NaH_2_PO_4_.2H_2_O pH 8, 300 mM NaCl, 50 mM Biotin. The protein was buffer exchanged into enterokinase cleavage buffer (20 mM Tris pH 8, 50 mM NaCl, 2 mM CaCl_2_) and cleaved using bovine enterokinase (EK, NEB) at 16U/mg protein for 4 hrs. The EK was removed using soybean trypsin inhibitor agarose (Sigma) and the cleaved NT* tag was removed using Ni-NTA resin. The purified HNoV NS1 protein was then buffer exchanged into 20 mM citrate phosphate buffer pH6.1, 150 mM NaCl and stored at −80°C.

### BJAB proliferation assay

10^6^ BJAB were washed with PBS-/-, resuspended in a PBS solution containing Cell Trace CFSE (Thermo Fisher C34554) and incubated for 20 min at 37°C. Next, cells were washed twice, resuspended in complete RPMI and treated or mock-treated with NS1 protein. At selected time points, cells were harvested in MagSep buffer, fixed/permeabilized with the Fixation/Permeabilization Solution Kit (BD Bioscience, 554714) and stained with the Alexa Fluor® 647-conjugate Ki-67 (D3B5) antibody (Cell Signaling, 12075S). Data was acquired with BD Fortessa and analyzed by using FlowJo. CFSE staining was measured by integrated intensity. The Ki-67 gate was defined on the non-treated BJAB to include <10% proliferating cells.

### Agilent Seahorse XF Real-Time ATP Rate Assay

The XF ATP Rate Assay (Agilent 103592-100) was used per the manufacturer’s instructions. Briefly, BJAB (5×10^4^ cells/well) were plated in 96 well plate in 80 mL of RPMI and treated with LPS (1ug/ml), NS1 protein (1ug/ml) or heat-inactivated NS1 protein (95°C for 10 minutes). 16 hrs after treatment, cells were transferred in a Cell Tak-coated Seahorse plate and allowed to adhere by centrifugation at 500 x g for 3 min. Cells were then washed 1X with 200 mL warm ATP Assay Medium prepared per protocol (Agilent XF DMEM Medium pH 7.4 (103757-100); 10 mM XF Glucose; 1 mM XF Sodium Pyruvate; 2 mM XF L-Glutamine) and then fresh ATP Assay medium was added for a final well volume of 180 mL. Cells were incubated at 37°C (non-CO_2_) for 30 minutes and ATP rate was quantified by using the Seahorse XFe96 Extracellular Flux Analyzer. Data were analyzed in the Agilent ATP Assay Report Generator and statistics were analyzed in Prism 7.0.

### Metabolomics analysis of intracellular metabolites

BJAB (10^6^ cells/well) were plated in 1ml of RPMI in 12-well plates and treated with LPS (1 μg/ml) or NS1 (1 µg/ml) for 16 hrs at 37°C. After incubation, BJAB were collected by centrifugation at 500 x g for 5 minutes at 4°C. The cell pellets were resuspended in ice-cold 80% methanol and kept at −80°C for 10 minutes. Supernatants were collected following centrifuge at the highest speed for 5 minutes at 4°C. Metabolites were dried at 4°C using a speedvac. Metabolite pellets were re-constituted in 50uL of 50% methanol and 40 µL was transferred to an auto-sampler glass vial for untargeted LC-MS analysis. Samples were run on an Agilent 1290 Infinity II LC −6470 Triple Quadrupole (QqQ) tandem mass spectrometer (MS/MS) system with the following parameters: Agilent Technologies Triple Quad 6470 LC-MS/MS system consists of the 1290 Infinity II LC Flexible Pump (Quaternary Pump), the 1290 Infinity II Multisampler, the 1290 Infinity II Multicolumn Thermostat with 6 port valve and the 6470 triple quad mass spectrometer. Agilent Masshunter Workstation Software LC/MS Data Acquisition for 6400 Series Triple Quadrupole MS with Version B.08.02 is used for compound optimization, calibration, and data acquisition.

#### LC

2uL of sample was injected into an Agilent ZORBAX RRHD Extend-C18 column (2.1 × 150 mm, 1.8 um) with ZORBAX Extend Fast Guards. The LC gradient profile is as follows, solvent conditions below. 0.25 ml/min, 0-2.5 min, 100% A; 2.5-7.5 min, 80% A and 20% B; 7.5min-13 min 55% A and 45% B; 13min-24 min, 1% A and 99% B; 24min-27min, 1% A and 99% C; 27min-27.5min, 1% A and 99% C; at 0.8 ml/min, 27.5-31.5 min, 1% A and 99% C; at 0.6 ml/min, 31.5-32.25min, 1% A and 99% C; at 0.4 ml/min, 32.25-39.9 min, 100% A; at 0.25 ml/min, 40 min, 100% A. Column temp is kept at 35°C, samples are at 4°C.

#### Solvents

Solvent A is 97% water and 3% methanol 15 mM acetic acid and 10 mM tributylamine at pH of 5. Solvent B is 15 mM acetic acid and 10 mM tributylamine in methanol. Washing Solvent C is acetonitrile. LC system seal washing solvent 90% water and 10% isopropanol, needle wash solvent 75% methanol, 25% water. Solvents were purchased from the following vendors: GC-grade Tributylamine 99% (ACROS ORGANICS), LC/MS grade acetic acid Optima (Fisher Chemical), InfinityLab Deactivator additive, ESI –L Low concentration Tuning mix (Agilent Technologies), LC-MS grade solvents of water, and acetonitrile, methanol (Millipore), isopropanol (Fisher Chemical).

#### MS

6470 Triple Quad MS is calibrated with the Agilent ESI-L Low concentration Tuning mix. Source parameters: Gas temp 150°C, Gas flow 10 l/min, Nebulizer 45 psi, Sheath gas temp 325°C, Sheath gas flow 12 l/min, Capillary −2000 V, Delta EMV −200 V. Dynamic MRM scan type is used with 0.07 min peak width, acquisition time is 24 min. Delta retention time of plus and minus 1 min, fragmentor of 40 eV and cell accelerator of 5 eV are incorporated in the method.

#### Data Analysis

The MassHunter Metabolomics Dynamic MRM Database and Method was used for target identification. Key parameters of AJS ESI were: Gas Temp: 150°C, Gas Flow 13 l/min, Nebulizer 45 psi, Sheath Gas Temp 325°C, Sheath Gas Flow 12 l/min, Capillary 2000 V, Nozzle 500 V. Detector Delta EMV(-) 200. The QqQ data were pre-processed with Agilent MassHunter Workstation QqQ Quantitative Analysis Software (B0700). Each metabolite abundance level in each sample was divided by the median of all abundance levels across all samples for proper comparisons, statistical analyses, and visualizations among metabolites.

## Acknowledgements

We thank Sasheen Dowlath for the technical support for the production of the NS1 recombinant protein. We further thank Dr. Jose G. Trevino and Dr. Steven Hughes, surgeons of the College of Medicine at the University of Florida, for performing surgical resections to obtain spleen and lymph node biopsies.

This work was supported by NIH grant R21AI130328 to CEW and R01AI123144 to SMK. CM was supported by MICHR training grant (LT1) and Marie-Slodowska Curie global fellowship. IAO was supported by a DELTAS Africa grant (DEL-15-007: Awandare). C.A.L. was supported by the NCI (R37CA237421). Metabolomics studies performed at the University of Michigan were supported by NIH grant DK097153, the Charles Woodson Research Fund, and the UM Pediatric Brain Tumor Initiative. MS was supported by the Kirschstein-NRSA F32 fellowship.

## Conflict of Interest

C.A.L. has received consulting fees from Astellas Pharmaceuticals and is an inventor on patents pertaining to Kras regulated metabolic pathways, redox control pathways in pancreatic cancer, and targeting the GOT1-pathway as a therapeutic approach.

